# Sequence Compression Benchmark (SCB) database — a comprehensive evaluation of reference-free compressors for FASTA-formatted sequences

**DOI:** 10.1101/642553

**Authors:** Kirill Kryukov, Mahoko Takahashi Ueda, So Nakagawa, Tadashi Imanishi

## Abstract

**Background:** Nearly all molecular sequence databases currently use gzip for data compression. Ongoing rapid accumulation of stored data calls for more efficient compression tool. Although numerous compressors exist, both specialized and general-purpose, choosing one of them was difficult because no comprehensive analysis of their comparative advantages for sequence compression was available.

**Findings:** We systematically benchmarked 410 settings of 44 compressors (including 26 specialized sequence compressors and 18 general-purpose compressors) on representative FASTA-formatted datasets of DNA, RNA and protein sequences. Each compressor was evaluated on 17 performance measures, including compression strength, as well as time and memory required for compression and decompression. We used 25 test datasets including individual genomes of various sizes, DNA and RNA datasets, and standard protein datasets. We summarized the results as the Sequence Compression Benchmark database (SCB database, http://kirr.dyndns.org/sequence-compression-benchmark/) that allows building custom visualizations for selected subsets of benchmark results.

**Conclusion:** We found that modern compressors offer large improvement in compactness and speed compared to gzip. Our benchmark allows comparing compressors and their settings using a variety of performance measures, offering the opportunity to select the optimal compressor based on the data type and usage scenario specific to particular application.

## Background

Molecular sequence databases store and distribute DNA, RNA and protein sequences as compressed FASTA-formatted files. Biological sequence compression was first proposed in 1986 [1] and the first practical compressor was made in 1993 [2]. A lively field emerged that produced a stream of methods, algorithms, and software tools for sequence compression [3,4]. However, despite this activity, currently nearly all databases universally depend on gzip for compressing FASTA-formatted sequence data. This incredible longevity of the 26-year-old compressor probably owes to multiple factors, including conservatism of database operators, wide availability of gzip, and its generally acceptable performance. Through all these years the amount of stored sequence data kept growing steadily [5], increasing the load on database operators, users, storage systems and network infrastructure. However, someone thinking to replace gzip invariably faces the questions: which of the numerous available compressors to choose? And will the resulting gains be even worth the trouble of switching?

Previous attempts at answering these questions are limited by testing too few compressors and by using restricted test data [6–11]. In addition, all of these studies provide results in form of tables, with no graphical outputs, which makes the interpretation difficult. Existing benchmarks with useful visualization such as Squash [12], are limited to general-purpose compressors.

The variety of available specialized and general-purpose compressors is overwhelming. At the same time the field was lacking a thorough investigation of comparative merits of these compressors for sequence data. Therefore we set out to benchmark all available and practically useful compressors on a variety of relevant sequence data. Specifically, we focused on the common task of compressing DNA, RNA and protein sequences, stored in FASTA format, without using reference sequence. The benchmark results were shown in the Sequence Compression Benchmark database (SCB database, http://kirr.dyndns.org/sequence-compression-benchmark/).

## Compressors and test data

We tested all sequence compressors that are available and functional in 2019: dnaX [13], XM [14], DELIMINATE [15], Pufferfish [16], DNA-COMPACT [17], MFCompress [18], UHT [19], GeCo [20], GeCo2 [21], JARVIS [22], NAF [23], and NUHT [24]. We also included the relatively compact among homology search database formats: BLAST [25] and 2bit - a database format of BLAT [26].

Since compressors designed for FASTQ data can be trivially adopted for FASTA-formatted inputs, we also included a comprehensive array of compressors designed primarily or specifically for FASTQ data: BEETL [27], Quip [28], fastqz [10], fqzcomp [10], DSRC 2 [29], Leon [30], LFQC [31], KIC [32], ALAPY [33], GTX.Zip [34], HARC [35], LFastqC [36], and SPRING [37].

We also tested a comprehensive array of general purpose compressors: bcm [38], brotli [39], bsc [40], bzip2 [41], cmix [42], gzip [43], lizard [44], lz4 [45], lzop [46], lzturbo [47], nakamichi [48], pbzip2 [49], pigz [50], snzip [51], xz [52], zpaq [53], zpipe [53] and zstd [54]. See Supplementary Table 1 for the list of compressors we used.

For the test data, we selected a variety of commonly used sequence datasets in FASTA format: (1) Individual genomes of various sizes, as examples of non-repetitive data; (2) DNA and RNA datasets, such as collections of mitochondrial genomes, influenza virus sequences, 16S rRNA gene sequences, and DNA alignments; (3) Standard protein datasets. Individual genomes are less repetitive, while other datasets are more repetitive. In total we used 25 test datasets. See Supplementary Table 2 for the list of test data.

## Benchmark

We benchmarked each compressor on every test dataset, except in cases of incompatibility (e.g., DNA compressors cannot compress protein data) or excessive time requirement (some compressors are so slow that they would take weeks on larger datasets). For compressors with adjustable compression level, we tested the relevant range of levels. We tested both 1 and 4-thread variants of compressors that support multithreading. In total, we used 410 settings of 44 compressors. We also included the non-compressing “cat” command as control. For compressors using wrappers, we also benchmarked the wrappers.

Currently many sequence analysis tools support gzip-compressed files as input. Switching to another compressor may require either adding support of new format to those tools, or passing the data in uncompressed form. The latter solution can be achieved with the help of Unix pipes, if both the compressor and the analysis tool support streaming mode. Therefore, we benchmarked all compressors in streaming mode (streaming uncompressed data in both compression and decompression).

For each combination of compressor setting and test dataset we recorded compressed size, compression time, decompression time, peak compression memory and peak decompression memory. The details of the method and raw benchmark data are available in Supplementary Methods and Supplementary Data, respectively. We share benchmark results and scripts via the SCM database at http://kirr.dyndns.org/sequence-compression-benchmark/.

The choice of measure for evaluating compressor performance depends on prospective application. For data archival, compactness is the single most important criterion. For public sequence database, the key measure is how long time it takes from initiating the download of compressed files until accessing the decompressed data. This time consists of transfer time plus decompression time (TD-Time). Corresponding transfer-decompression speed (TD-Speed) is computed as Original Size / TD-Time. In this use case compression time is relatively unimportant, since compression happens only once, while transfer and decompression times affect every user of the database. For one-time data transfer, all three steps of compression, transfer and decompression are timed (CTD-Time), and used for computing the resulting overall speed (CTD-Speed).

A total of 17 measures, including the above-mentioned ones, are available in our results data (See Supplementary Methods for the list of measures). Any of these measures can be used for selecting the best setting of each compressor and for sorting the list of compressors. These measures can be then shown in a table and visualized in column charts and scatterplots. This allows tailoring the output to answer specific questions, such as what compressor is better at compressing particular kind of data, or which setting of each compressor performs best at particular task. The link speed that is used for estimating transfer times is configurable. The default speed of 100 Mbit/sec corresponds to the common speed of a fixed broadband internet connection.

Fig.1 compares the performance of best settings of 35 compressors on human genome. It shows that specialized sequence compressors achieve excellent compression ratio on this genome. However, when total TD-Speed or CTD-Speed is considered (measures that are important in practical applications), most sequence compressors fall behind the general-purpose ones. The best compressors for this dataset in terms of compression ratio, TD-Speed and CTD-Speed are “fastqz-slow”, “naf-22” and “naf-1”, respectively (numbers in each compressor name indicate compression level and other settings). Interestingly, the non-compressing “cat” command used as control, while naturally showing at the last place on compression ratio (Fig.1A), is not the slowest in terms of TD-Speed and CTD-Speed (Figs.1B and 1C, respectively). In case of CTD-Speed, for example, it means that some compressors are so slow that their compression + transfer + decompression time turns out to be longer than time required for transferring raw uncompressed data (using particular link speed, in this case 100 Mbit/sec).

**Fig. 1.**
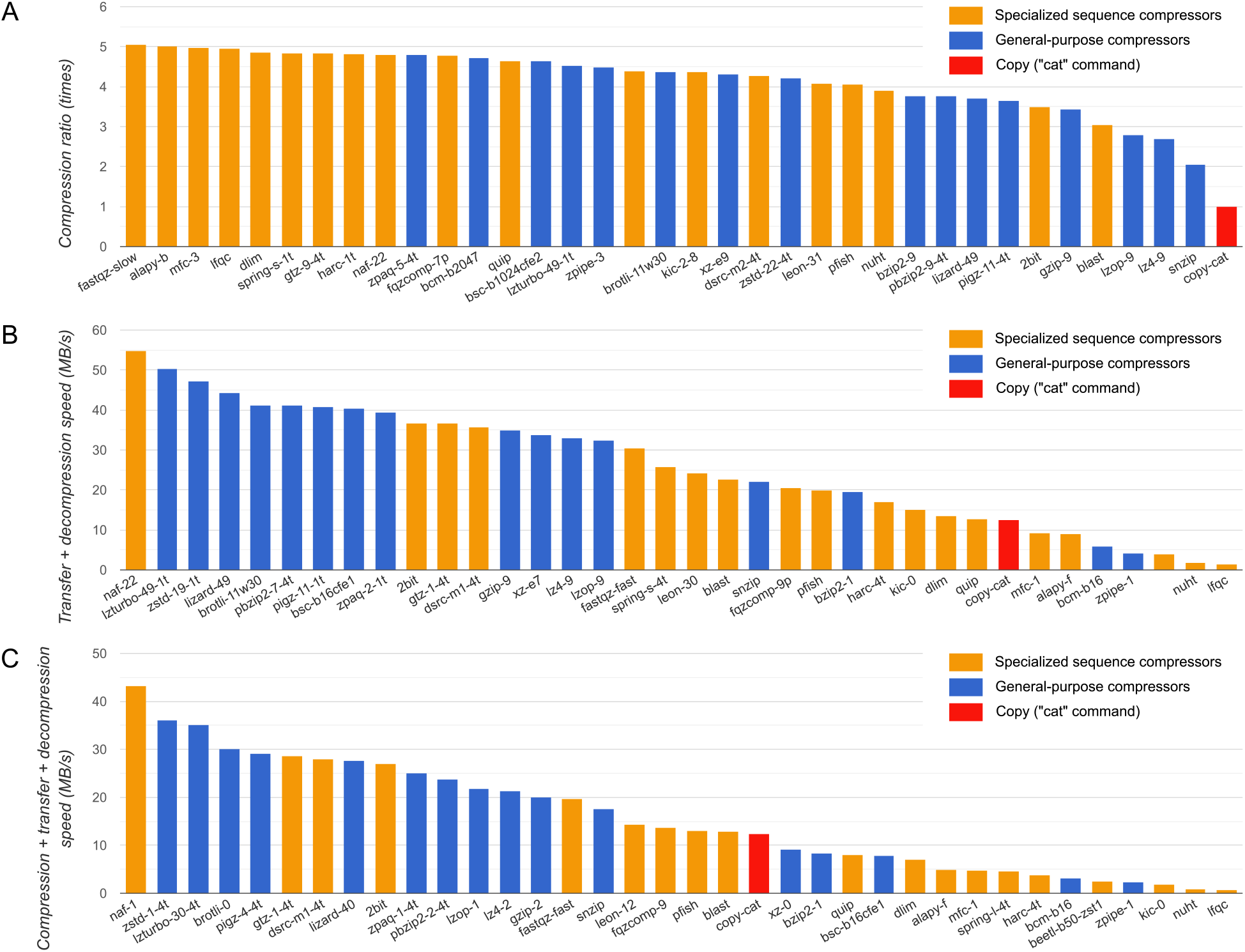
Comparison of 35 compressors on human genome. Best settings of each compressor are selected based on different aspects of performance: (A) compression ratio, (B) transfer + decompression speed, and (C) compression + transfer + decompression speed. Specialized sequence compressors are shown in orange color, and general-purpose compressors are shown in blue. The copy-compressor (“cat” command), shown in red color, is included as a control. The selected settings of each compressor are shown in their names, after hyphen. Multi-threaded compressors have “-1t” or “-4t” at the end of their names to indicate the number of threads used. Test data is the 3.31 GB reference human genome (accession number GCA_000001405.28). Benchmark CPU: Intel Xeon E5-2643v3 (3.4 GHz). Link speed of 100 Mbit/s was used for estimating the transfer time.

Fig.2 compares all compressor settings on the same data (human genome). Fig.2A shows that the strongest compressors often provide very slow decompression speed (shown using logarithmic scale due to the enormous range of values), which means that quick data transfer (resulting from strong compression) offered by those compressors is offset by significant waiting time required for decompressing the data. Fig.2B shows TD-Speed plotted against CTD-Speed. Similar figures can be constructed for other data and performance measures on the SCB database website.

**Fig. 2.**
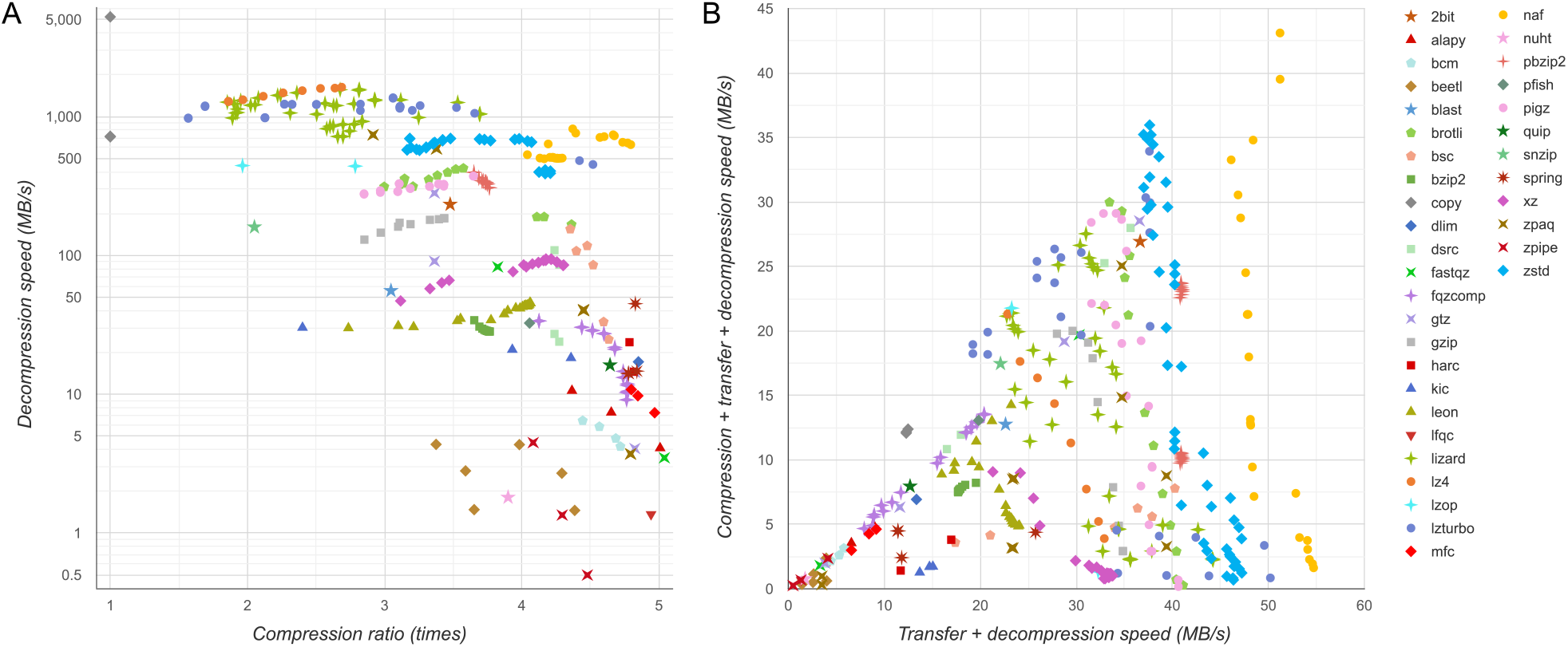
Comparison of 335 settings of 36 compressors on human genome. Each point represents a particular setting of some compressor. Panel A shows the relationship between compression ratio and decompression speed. Panel B shows the transfer + decompression speed plotted against compression + transfer + decompression speed. Test data is the 3.31 GB reference human genome (accession number GCA_000001405.28). Benchmark CPU: Intel Xeon E5-2643v3 (3.4 GHz). Link speed of 100 Mbit/s was used for estimating the transfer time.

Visualizing results from multiple test datasets simultaneously is possible, with or without aggregation of data. With aggregation, the numbers will be summed or averaged, and a single measurement will be shown for each setting of each compressor. Without aggregation, the results of each compressor setting will be shown separately on each dataset. Since the resulting number of data points can be huge, in such case it is useful to request only best settings of each compressor to be shown. The criteria for choosing the best setting is selectable among the 17 measurements. In case of a column chart, any of the 17 measures can be used for ordering the shown compressors, independently of the setting used for selecting best version, and independently for the measure actually shown in the chart.

One useful capability of the SCB database is showing measurements relative to specified compressor (and setting). This allows selecting a reference compressor and comparing the other compressors to this reference. For example, we can compare compressors to gzip as shown on Fig.3. In this example, we compare only best settings of each compressor, selected using specific measures (transfer+decompression speed and compression+transfer+decompression speed on Figs.3A and 3B, respectively). We also used fixed scale to show only range above 0.5 on both axes, which means that only performances that are at least half as good as gzip on both axes as shown. In this example, we can see that some compressors improve compactness and some improve speed compared to gzip, but few compressors improve both at the same time, such as lizard, naf, pigz, pbzip, and zstd.

**Fig. 3.**
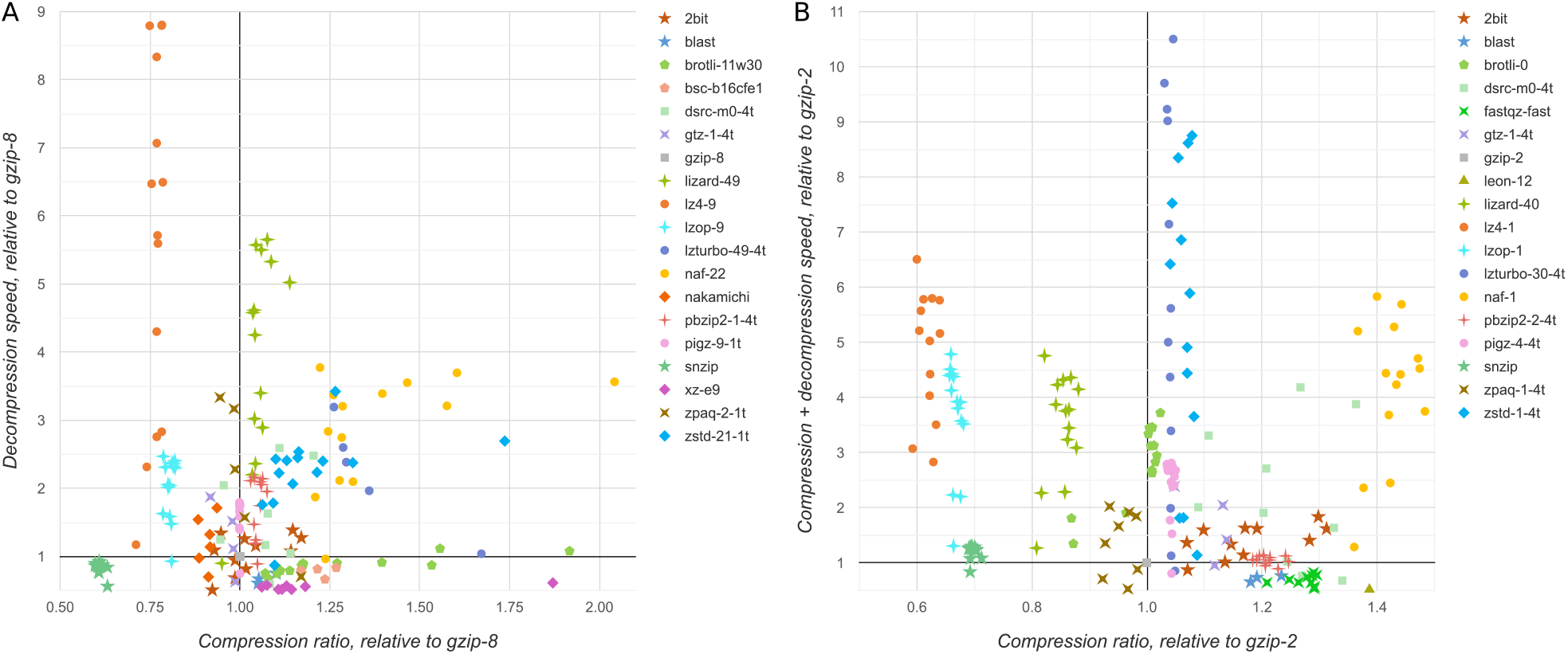
Comparison of compressor settings to gzip. Genome datasets were used as test data. Each point shows the performance of a compressor setting on specific genome test dataset. All values are shown relative to representative setting of gzip. Only performances that are at least half as good as gzip on both axes are shown. Panel A shows settings that performed best in Transfer+Decompression speed, B - settings that performed best in Compression+Transfer+Decompression speed. Link speed of 100 Mbit/s was used for estimating the transfer time. The grid lines crossing the (1,1) coordinate are highlighted.

It is important to be aware of the memory requirements when choosing a compressor (Fig.4). In these charts we plotted data size on the X axis, and disabled aggregation. This allows seeing how much memory a particular compressor used on each test dataset. As this example shows memory requirement reaches saturation point for most compressors. On the other hand, some compressors have unbounded growth of consumed memory, which makes then unusable for large data. Interestingly, gzip apparently has the smallest memory footprint, which may be one of the reasons for its popularity. Most compressors can function on a typical desktop hardware, but some require larger memory, which is important to consider when choosing a compressor that will be run by the consumers of distributed data.

**Fig. 4.**
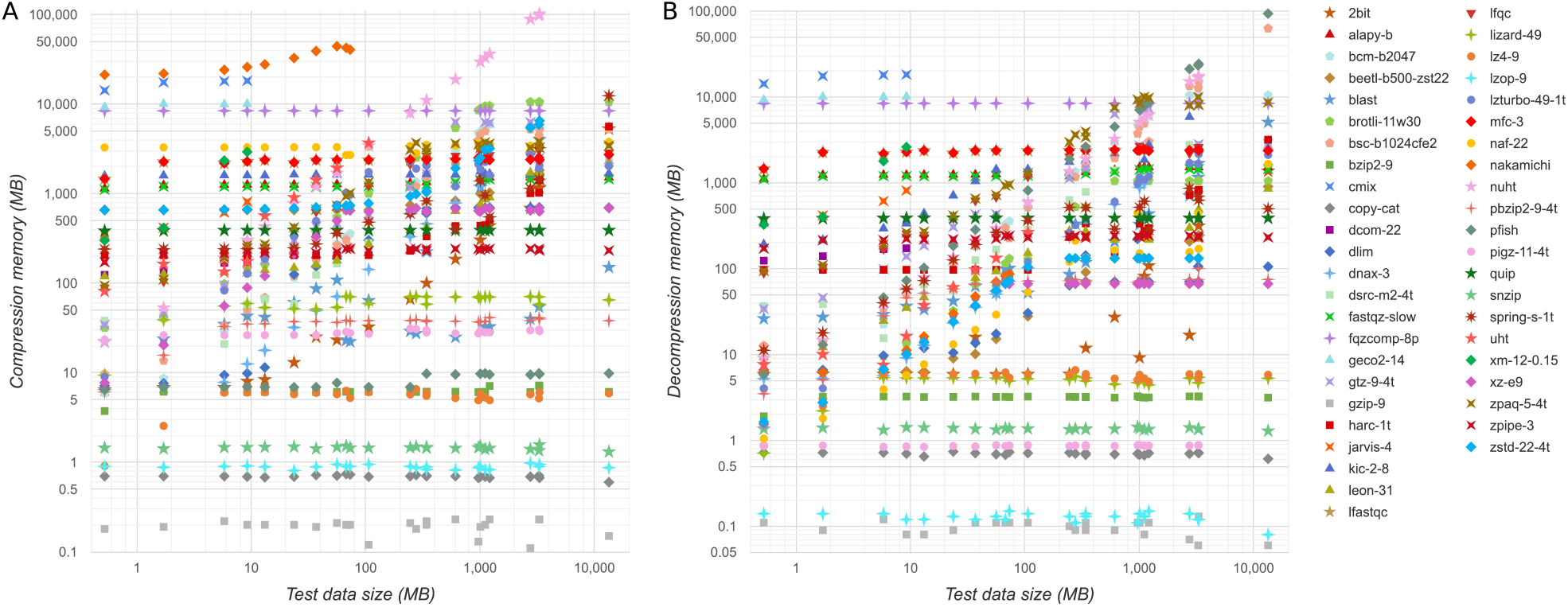
Compressor memory consumption. Strongest setting of each compressor is shown. On the X axis is the test data size. On the Y axis is the peak memory used by the compressor, for compression (A) and decompression (B).

Wide variety of charts can be produced on the benchmark website by selecting specific combinations of test data, compressors, and performance measures. At any point the currently visualized data can be obtained in textual form using Table output option. Also, all charts can be downloaded in SVG format.

## Conclusions

Our benchmark reveals complex relationship between compressors and between their settings, based on various measures. We found that continued use of gzip is usually far from an optimal choice. Transitioning from gzip to a better compressor brings significant gains for genome and protein data, and is especially beneficial with repetitive DNA/RNA datasets. Overall, our data suggests using naf-22 as the default compressor to archive FASTA-formatted sequences, because it combines good compression strength with very quick decompression. However, it is best to check the results for specific data types and performance measures.

The Sequence Compression Benchmark (SCB) database will help in navigating the complex landscape of data compression. With dozens of compressors available, making an informed choice is not an easy task and requires careful analysis of the project requirements, data type and compressor capabilities. Our benchmark is the first resource providing a detailed practical evaluation of various compressors on a wide range of molecular sequence datasets. Using the SCB database, users can analyze compressor performances on variety of metrics, and construct custom reports for answering project-specific questions.

In contrast to previous studies that showed their results in static tables, our project is dynamic in two important senses: (1) the result tables and charts can be dynamically constructed for a custom selection of test data, compressors, and measured performance numbers, and (2) our study is not a one-off benchmark, but marks the start of a project where we will continue to add compressors and test data.

Making an informed choice of compressor with the help of our benchmark will lead to increased compactness of sequence databases, with shorter time required for downloading and decompressing. This will reduce the load on network and storage infrastructure, and increase speed and efficiency in biological and medical research.

## Declarations

### Availability of data and material

All benchmark data is available at the online SCB database: http://kirr.dyndns.org/sequence-compression-benchmark/

### Competing interests

The authors declare no competing interests.

### Funding

This work was supported by the 2019 Tokai University School of Medicine Research Aid (to KK), JSPS KAKENHI Grants-in-Aid for Scientific Research on Innovative Areas (16H06429, 16K21723, 19H04843 to SN), and Takeda Science Foundation (to TI).

### Authors’ contributions

KK conceived the study idea and implemented the benchmark. SN provided benchmark hardware. KK, MTU, SN and TI interpreted the data and wrote the manuscript. KK and MTU prepared figures and tables. All authors read and approved the final manuscript.

## Acknowledgements

Not applicable.

---

**Supplementary Table 1.**
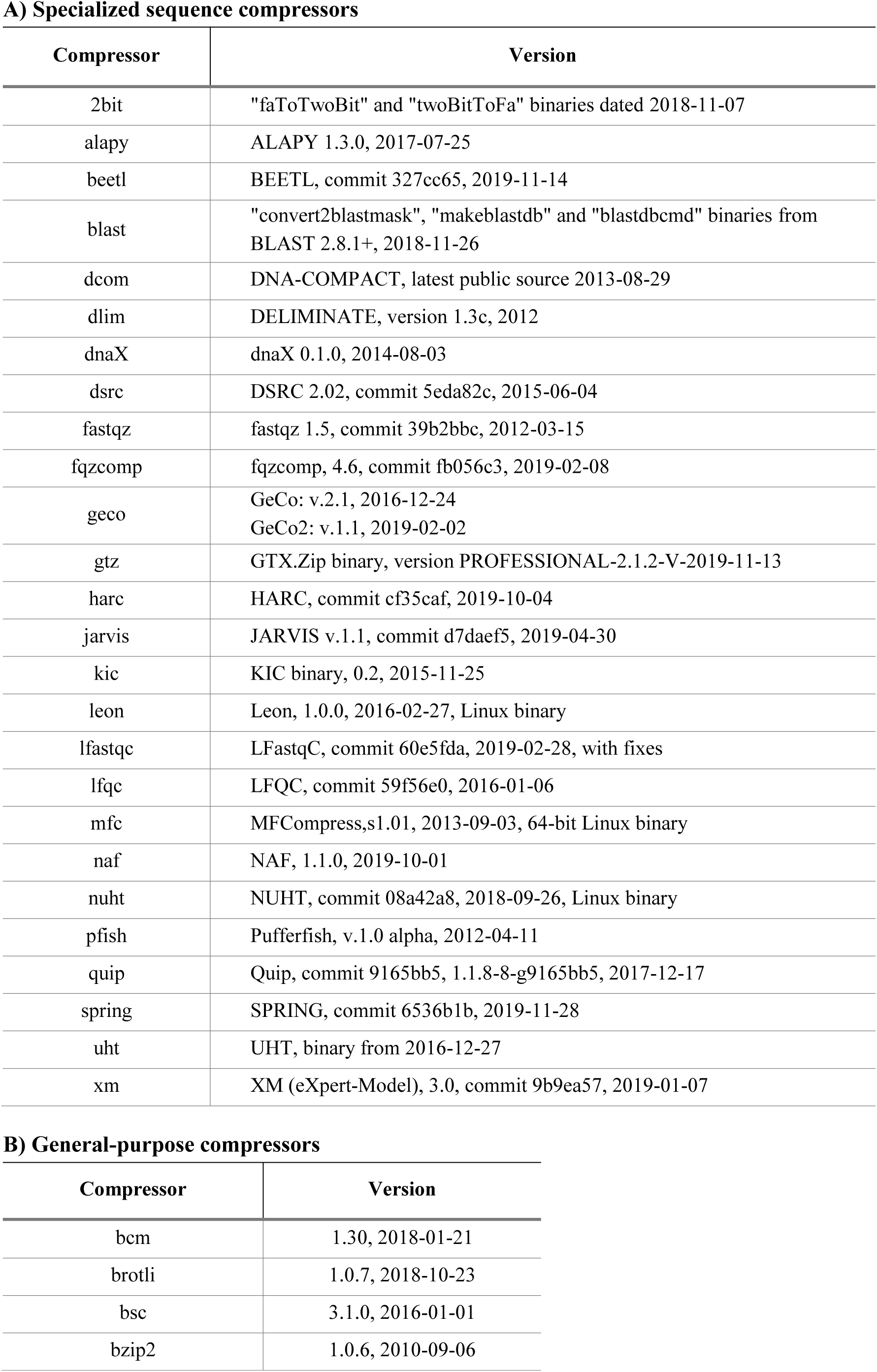
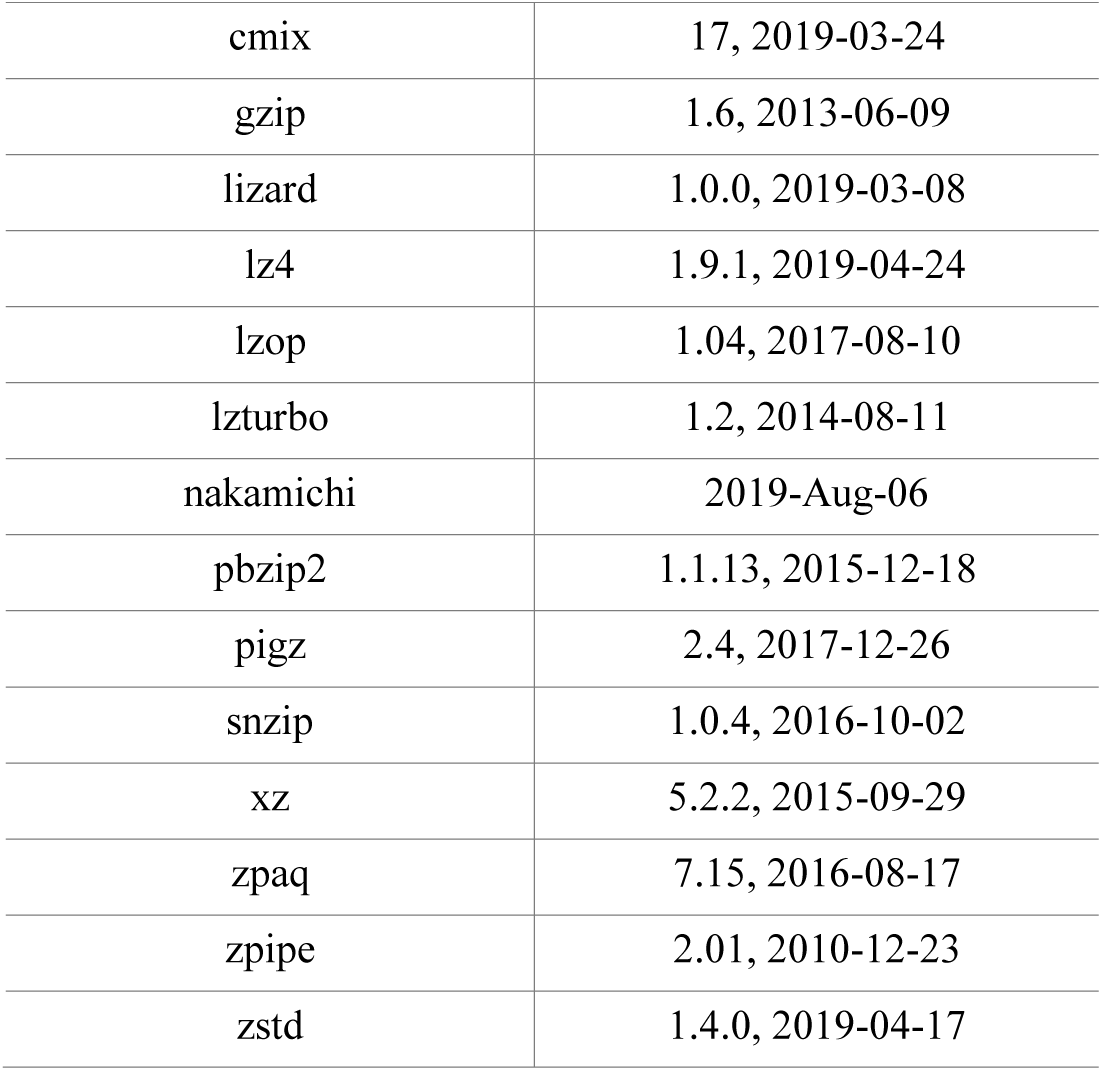
Compressor versions

**Supplementary Table 2.**
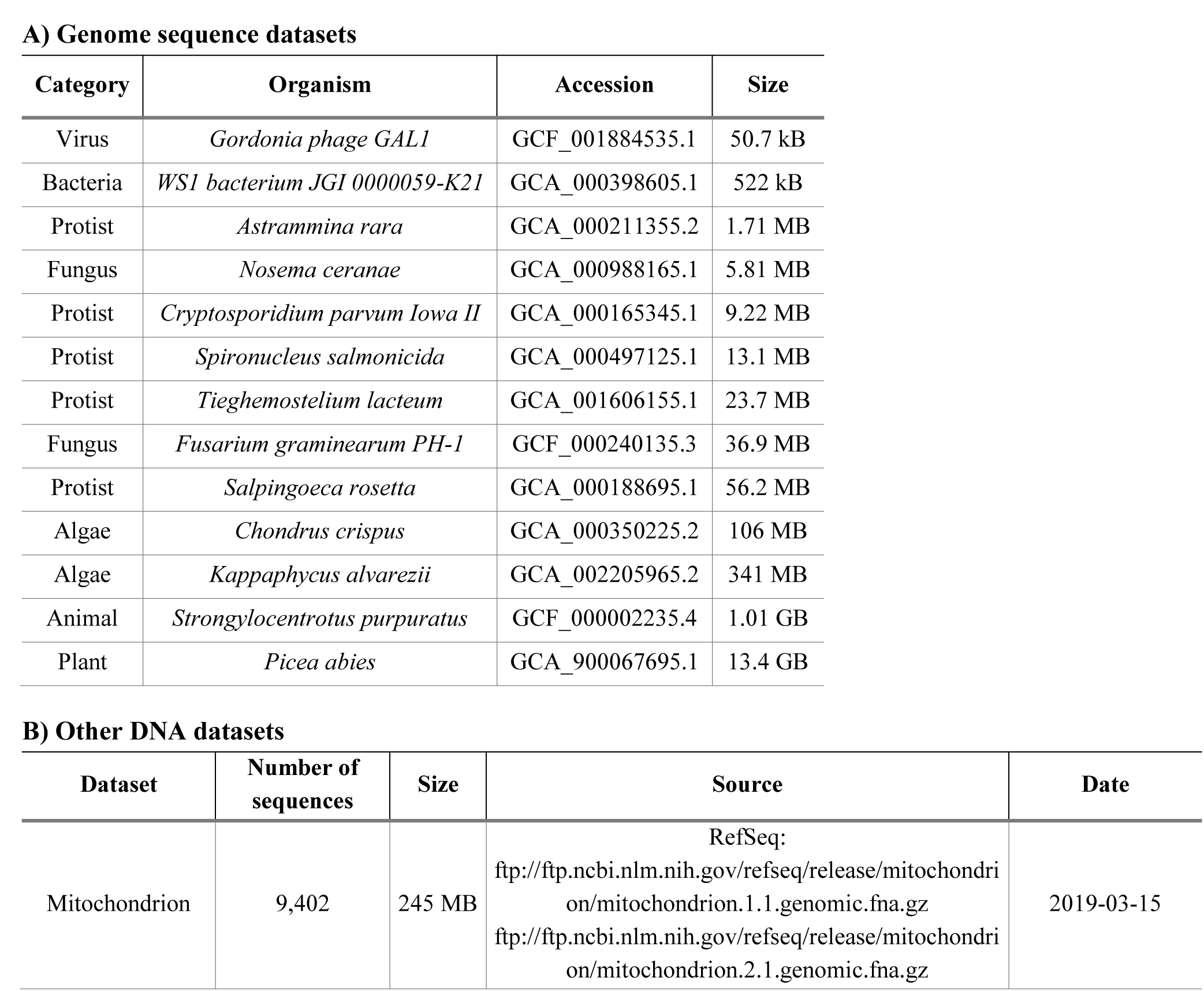
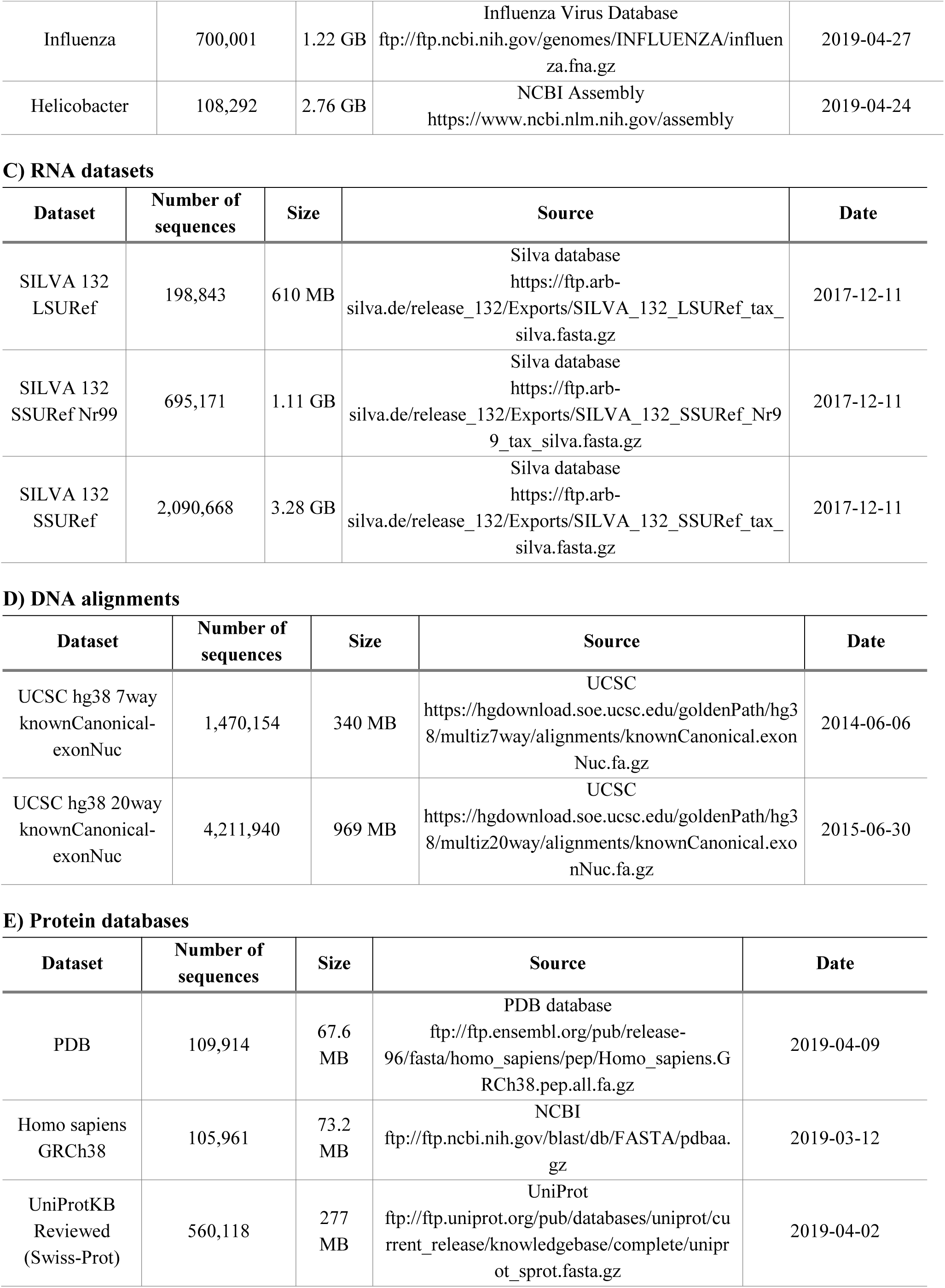
Test datasets

## Supplementary Methods

### Compression task

The task is to compress and decompress a FASTA-formatted file containing DNA, RNA or protein sequences. The process should be lossless, i.e., decompressed data should be identical to the original data. Compression and decompression are done without using any reference genome. Each compression and decompression runs in Linux in a command line interface. Input data for compression and output data during decompression are streamed using Unix pipes.

### Compressor selection

We used all specialized sequence compressors that we could find and make to work for the above specified task. For general-purpose compressors we used only the major ones, in terms of performance, historical importance, or popularity. For each compressor with configurable compression level (or other parameters related to compression strength of speed), we used the relevant range of settings, including the default.

### Benchmark machine

- CPU: dual Xeon E5-2643v3 (3.4 GHz, 6 cores), hyperthreading: off
- RAM: 128 GB DDR4-2133 ECC Registered
- Storage: 4 x 2 TB SSD, in RAID 0, XFS filesystem, block size: 4096 bytes (blockdev--getbsz)
- OS: Ubuntu 18.04.1 LTS, kernel: 4.15.0
- GCC: 7.4.0

### Compressor/dataset combinations that were tested

Each setting of each compressor is tested on every test dataset, except when it’s difficult or impossible due to compressor limitations:

- Due to their extreme slowness, these compressors are not tested on any data larger than 10 MB: cmix, DNA-COMPACT, GeCo, JARVIS, Leon, and XM.
- UHT fails on the 245 MB dataset and on larger data.
- Nakamichi was only used on data smaller than 100 MB due to its slowness and memory resultements.
- Among sequence compressors, only DELIMINATE, MFCompress and NAF support alignments.
- Among sequence compressors, only BLAST and NAF support protein sequences.
- Some settings of XM crash and/or produce wrong decompressed output on some data - such results are not included.
- NUHT’s memory requirement makes it impossible to use on 13.4 GB *Picea abies* genome.

### Benchmark process

The entire benchmark is orchestrated by a perl script. This script loads the lists of compressor settings and test data, and proceeds to test each combination that still has its measurements missing in the output directory. For each such combination (of compressor setting and test dataset), the following steps are performed:

1. Compression is performed by piping the test data into the compressor. Compressed size and compression time is recorded. For compressed formats consisting of multiple files, sizes of all files are summed together.
2. If compression time did not exceed 10 seconds, 9 more compression runs are performed, recording compression times. Compressed data from previous run is deleted before each next compression run.
3. The next set of compression runs is performed to measure peak memory consumption. This set consists of the same number of runs as in steps 1-2 (either 1 or 10 runs). That is, for fast compressors and for small data the measurement is repeated 10 times.
4. Decompression test run is performed. In this run decompressed data is piped to the “md5sum -b -” command. The resulting md5 signature is compared with that from the original file. In case of any mismatch this combination of compressor setting and dataset is disqualified and its measurements are discarded.
5. Decompression time is measured. This time decompressed data is piped to /dev/null.
6. If decompression completed within 10 seconds, 9 more decompression runs are performed and timed.
7. Peak decompression memory is measured. The number of runs is same as in steps 5-6.
8. The measurements are stored to a file. All compressed and temporary files are removed.

### Measurement methods

Measuring time: Wall clock time was measured using Perl’s Time::HiRes module (gettimeofday and tv_interval subroutines). The resulting time was recorded with millisecond precision.

Measuring peak memory consumption: First, each compression command was stored in a temporary shell script file. Then it was executed via GNU Time, as /usr/bin/time -v cmd.sh >output.txt. “Maximum resident set size” value was extracted from the output. 1638 was then subtracted from this value and the result was stored as peak memory measurement. 1638 is the average “Maximum resident set size” measured by GNU Time in the same way for an empty script.

Memory consumption and time were measured separately because measuring memory makes the task noticeably slower, especially for very fast tasks.

### Collected measurements

For each combination of compressor and dataset that was tested, the following measurements were collected:

- Compressed size (in bytes)
- Compression time (in milliseconds)
- Decompression time (in milliseconds)
- Peak compression memory (in GNU Time’s “Kbytes”)
- Peak decompression memory (in GNU Time’s “Kbytes”)

In cases where 10 values are collected, the average value is used by the benchmark web-site.

### Computed values

The following values were calculated based on the measured values:

- Compressed size relative to original (%) = Compressed size / Uncompressed size * 100
- Compression ratio (times) = Uncompressed size / Compressed size
- Compression speed (MB/s) = Uncompressed size in MB / Compression time
- Decompression speed (MB/s) = Uncompressed size in MB / Decompression time
- Compression + decompression time (s) = Compression time + Decompression time
- Compression + decompression speed (MB/s) = Uncompressed size in MB / (Compression time + Decompression time)
- Transfer time (s) = Uncompressed size / Link speed in B/s
- Transfer speed (MB/s) = Uncompressed size in MB / Transfer time
- Transfer + decompression time (s) = Transfer time + Decompression time
- Transfer + decompression speed (MB/s) = Uncompressed size in MB / (Transfer time + Decompression time)
- Compression + transfer + decompression time (s) = Compression time + Transfer time + Decompression time
- Compression + transfer + decompression speed (MB/s) = Uncompressed size in MB / (Compression time + Transfer time + Decompression time)

### Rationale for non-constant number of runs

Variable number of runs is the only way to have both accurate measurements and large test data (under the constraints of using one test machine, and running benchmark within reasonable time).

On one hand, benchmark takes lot of time. So much that some compressors can’t be even tested at all on dataset larger than 10 MB in reasonable time. Therefore repeating every measurement 10 times is impractical. Or, it would imply restricting the test data to only small datasets.

On the other hand, measurements are slightly noisy. The shorter measured time, the more noisy its measurement. Thus for very quick runs, multiple runs allow for substantial noise suppression. For longer runs it does not make much difference, because the relative error is already small with longer times.

Using a threshold of 10 seconds seems a reasonable compromise between suppressing noise and including larger test data (and slow compressors).

### Compressor setup

For compression, each compressor was reading the input data streamed via unix pipe (“|” in the command line). For decompression, each compressor was set up to stream the decompressed data via pipe. This was done to better approximate the common pattern of using a compressor in a practical application. In an actual sequence analysis workflow, often the decompressed data is piped directly into the downstream analysis command. Also, when compressing the sequences, often the data is first pre-processed with another command, which then pipes the processed sequences to the compressor.

### Wrappers

Since some compressors do not support such streaming mode of operation, we used them via wrapper scripts. Our wrapper scripts also work around other deficiencies of many compressors, and add the following features to the compressors missing them:

- Supporting RNA input for DNA-only compressors.
- Supporting ‘N’ in DNA/RNA sequences.
- Supporting IUPAC’s ambiguous nucleotide codes.
- Saving and restoring line lengths.
- Saving and restoring sequence names.
- Saving and restoring sequence mask (upper/lower case).
- Supporting FASTA-formatted input.
- Supporting input with more than 1 sequence.

All our wrappers and commands are available at the SCB database website (http://kirr.dyndns.org/sequence-compression-benchmark/).

## Notes

#### Summary of Updates

Added test data and compressors, improved methodology, updated results and discussion.

http://kirr.dyndns.org/sequence-compression-benchmark/

## References

1. Walker JR, Willett P. Compression of nucleic acid and protein sequence data. Comput. Appl. Biosci. 1986;2(2): 89–93.

2. Grumbach S, Tahi F. Compression of DNA sequences. Data Compression Conference, Snowbird, Utah, IEEE Computer Society. 1993. p. 340–50. doi:10.1109/DCC.1993.253115.

3. Deorowicz S, Grabowski S. Data compression for sequencing data. Algorithms for Molecular Biology. 2013;8:25. doi:10.1186/1748-7188-8-25.

4. Hernaez M, Pavlichin D, Weissman T, Ochoa I. Genomic Data Compression. Annual Review of Biomedical Data Science. 2019;2:19–37. doi:10.1146/annurev-biodatasci-072018-021229.

5. Karsch-Mizrachi I, Takagi T, Cochrane G. The international nucleotide sequence database collaboration. Nucleic Acids Res. 2018;46(Database issue):D48–D51. doi:10.1093/nar/gkx1097.

6. Zhu Z, Zhang Y, Ji Z, He S, Yang X. High-throughput DNA sequence data compression. Brief. Bioinform. 2013; 16(1):1–15. doi:10.1093/bib/bbt087.

7. Hosseini M, Pratas D, Pinho AJ. A Survey on Data Compression Methods for Biological Sequences. Information. 2016;7(4):56. doi:10.3390/info7040056.

8. Sardaraz M, Tahir M. Advances in high throughput DNA sequence data compression. J. Bioinform. Comput. Biol. 2016;14(3):1630002. doi:10.1142/S0219720016300021.

9. Biji CL, Achuthsankar SN. Benchmark Dataset for Whole Genome Sequence Compression. IEEE/ACM Trans. Comput. Biol. Bioinform. 2017;14(6):1228–36. doi:10.1109/TCBB.2016.2568186.

10. Bonfield JK, Mahoney MV. Compression of FASTQ and SAM Format Sequencing Data. PLoS One. 2013;8(3): e59190, doi:10.1371/journal.pone.0059190.

11. Numanagic I, Bonfield JK, Hach F, Voges J, Ostermann J, Alberti C, Mattavelli M. Comparison of high-throughput sequencing data compression tools. Nature Methods. 2016;13(12):1005–8, doi:10.1038/nmeth.4037.

12. Squash Compression Benchmark. 2015. https://quixdb.github.io/squash-benchmark/. Accessed July 15, 2019.

13. Manzini G, Rastero M. A simple and fast DNA compressor. Software - Practice and Experience. 2004;34:1397–411, doi:10.1002/spe.619.

14. Cao MD, Dix TI, Allison L. Mears C. A simple statistical algorithm for biological sequence compression. Data Compression Conference. DCC ‘07, Snowbird, UT, IEEE Computer Society. 2007. p. 43–52. doi:10.1109/DCC.2007.7.

15. Mohammed MH, Dutta A, Bose T, Chadaram S, Mande SS. DELIMINATE — a fast and efficient method for loss-less compression of genomic sequences. Bioinformatics. 2012;28:2527–29. doi:10.1093/bioinformatics/bts467.

16. Pufferfish. 2012. https://github.com/alexholehouse/pufferfish. Accessed May 23, 2019.

17. Li P, Wang S, Kim J, Xiong H, Ohno-Machado L, Jiang X. DNA-COMPACT: DNA COMpression Based on a Pattern-Aware Contextual Modeling Technique. PLoS ONE. 2013;8(11):e80377. doi:10.1371/journal.pone.0080377.

18. Pinho AJ, Pratas D. MFCompress: a compression tool for FASTA and multi-FASTA data. Bioinformatics. 2014;30:117–8. doi:10.1093/bioinformatics/btt594.

19. Al-Okaily A, Almarri B, Al Yami S, Huang CHs. Toward a Better Compression for DNA Sequences Using Huffman Encoding. J. Comp. Biol. 2017;24(4):280–8. doi:10.1089/cmb.2016.0151.

20. Pratas D, Pinho AJ, Ferreira PJSG. Efficient compression of genomic sequences. *Data Compression Conference*, DCC-2016, Snowbird, Utah, IEEE Computer Society. 2016. p.231–240. doi: 10.1109/DCC.2016.60.

21. Pratas D, Hosseini M, Pinho AJ. GeCo2: An Optimized Tool for Lossless Compression and Analysis of DNA Sequences. *Practical Applications of Computational Biology and Bioinformatics, 13th International Conference*, PACBB 2019, Advances in Intelligent Systems and Computing, vol 1005, Springer, Cham, 2019a. p.137–145. doi: 10.1007/978-3-030-23873-5_17.

22. Pratas D, Pratas D, Hosseini M, Silva J, Pinho AJ. A Reference-Free Lossless Compression Algorithm for DNA Sequences Using a Competitive Prediction of Two Classes of Weighted Models. Entropy, 2019b;21:1074. doi: 10.3390/e21111074.

23. Kryukov K, Ueda MT, Nakagawa S, Imanishi T. Nucleotide Archival Format (NAF) enables efficient lossless reference-free compression of DNA sequences. Bioinformatics. 2019;35(19):3826–28. doi: 10.1093/bioinformatics/btz144.

24. Alyami, S, Huang CH. Nongreedy Unbalanced Huffman Tree Compressor for Single and Multifasta Files. Journal of Computational Biology. 2019; 26(0):1–9. doi: 10.1089/cmb.2019.0249.

25. Altschul SF, Gish W, Miller W, Myers EW, Lipman DJ. Basic local alignment search tool. J. Mol. Biol. 1990;215(3):403–10. doi: 10.1016/S0022-2836(05)80360-2.

26. Kent WJ. BLAT - The BLAST-Like Alignment Tool. Genome Research. 2002;12(4):656–64. doi: 10.1101/gr.229202.

27. Bauer MJ, Cox AJ, Rosone G. Lightweight BWT Construction for Very Large String Collections. *Combinatorial Pattern Matching 2011*, proceedings of the CPM 2011, 2011. p.219–231. doi: 10.1007/978-3-642-21458-5_20.

28. Jones DC, Ruzzo WL, Peng X, Katze MG. Compression of next-generation sequencing reads aided by highly efficient de novo assembly. Nucleic Acids Research. 2012;40(22):e171. doi: 10.1093/nar/gks754.

29. Roguski L, Deorowicz S. DSRC 2—Industry-oriented compression of FASTQ files. Bioinformatics. 2014; 30(15):2213–5. doi: 10.1093/bioinformatics/btu208.

30. Benoit G, Lemaitre C, Lavenier D, Drezen E, Dayris T, Uricaru R6, Rizk G. Reference-free compression of high throughput sequencing data with a probabilistic de Bruijn graph. BMC Bioinformatics. 2015;16:288. doi: 10.1186/s12859-015-0709-7.

31. Nicolae M, Pathak S, Rajasekaran S. LFQC: a lossless compression algorithm for FASTQ files. Bioinformatics. 2015;31(20):3276–81. doi: 10.1093/bioinformatics/btv384.

32. Zhang Y, Patel K, Endrawis T, Bowers A4, Sun Y. A FASTQ compressor based on integer-mapped k-mer indexing for biologist. Gene. 2016;579(1):75–81. doi: 10.1016/j.gene.2015.12.053.

33. ALAPY 2017. http://alapy.com/services/alapy-compressor/. Accessed December 2, 2019.

34. Xing Y, Li G, Wang Z, Feng B, Song Z, Wu C. GTZ: a fast compression and cloud transmission tool optimized for FASTQ files. BMC Bioinformatics. 2017;18(Suppl 16):549. doi: 10.1186/s12859-017-1973-5.

35. Chandak S, Tatwawadi K, Weissman T. Compression of genomic sequencing reads via hash-based reordering: algorithm and analysis. Bioinformatics. 2018; 34(4):558–67. doi: 10.1093/bioinformatics/btx639.

36. Al Yami S, Huang CH. LFastqC: A lossless non-reference-based FASTQ compressor. PLoS One. 2019;14(11):e0224806, doi: 10.1371/journal.pone.0224806.

37. Chandak S, Tatwawadi K, Ochoa I, Hernaez M, Weissman T. SPRING: a next-generation compressor for FASTQ data. Bioinformatics. 2019;35(15):2674–6. doi: 10.1093/bioinformatics/bty1015.

38. BCM. 2018. https://github.com/encode84/bcm. Accessed June 6 2019.

39. Alakuijala J, Szabadka Z. Brotli Compressed Data Format. RFC 7932. 2016. Accessed April 14 2019.

40. libbsc. 2016. https://github.com/IlyaGrebnov/libbsc. Accessed June 22 2019.

41. bzip2. 2019. https://www.sourceware.org/bzip2/. Accessed January 20 2019.

42. cmix. 2019. https://github.com/byronknoll/cmix. Accessed April 25 2019.

43. GNU Gzip. 2018. https://www.gnu.org/software/gzip/. Accessed November 8 2019.

44. Lizard - efficient compression with very fast decompression. 2019. https://github.com/inikep/lizard. Accessed June 16 2019.

45. LZ4 - Extremely fast compression. 2019. https://github.com/lz4/lz4. Accessed April 25 2019.

46. Lzop. 2017. https://www.lzop.org/. Accessed December 6 2018.

47. LzTurbo - World’s fastest compressor. 2018. https://sites.google.com/site/powturbo/. Accessed February 11 2019.

48. Nakamichi. 2019. http://www.sanmayce.com/Nakamichi/index.html. Accessed December 1 2019.

49. pbzip2. 2015. https://launchpad.net/pbzip2/. Accessed April 26 2019.

50. pigz. 2017. https://zlib.net/pigz/. Accessed April 26 2019.

51. Snzip, a compression/decompression tool based on snappy. 2016. https://github.com/kubo/snzip. Accessed November 11 2018.

52. XZ Utils. 2018. https://tukaani.org/xz/. Accessed December 17 2018.

53. ZPAQ Incremental Journaling Backup Utility and Archiver. 2016. http://www.mattmahoney.net/dc/zpaq.html. Accessed November 7 2018.

54. Zstandard - Fast real-time compression algorithm. 2019. https://github.com/facebook/zstd. Accessed April 25 2019.

